# Development of a whole cell biosensor for detection of 2, 4-diacetylphloroglucinol-producing bacteria from grassland soil

**DOI:** 10.1101/2020.06.23.168377

**Authors:** Morten Lindqvist Hansen, Zhiming He, Mario Wibowo, Lars Jelsbak

## Abstract

Fluorescent *Pseudomonas* spp. producing the antibiotic 2,4-diacetylphloroglucinol (DAPG) are ecologically important in the rhizosphere as they can control phytopathogens and contribute to disease suppressiveness. While studies of DAPG-producing *Pseudomonas* have predominantly focused on rhizosphere niches, the ecological role of DAPG as well as the distribution and dynamics of DAPG-producing bacteria remains less well understood for other environments such as bulk soil and grassland, where the level of DAPG producers are predicted to be low. Here, we construct a whole cell biosensor for detection of DAPG and DAPG-producing bacteria from environmental samples.

We show that the sensor is highly specific towards DAPG, with a sensitivity in the low nanomolar range (<20 nm). This sensitivity is comparable to the DAPG levels identified in rhizosphere samples by chemical analysis. The biosensor enables guided isolation of DAPG-producing *Pseudomonas*. Using the biosensor, we probed the same grassland soil sampling site to isolate genetically related DAPG-producing *Pseudomonas kilonensis* strains over a period of 12 months. Next, we used the biosensor to determine the frequency of DAPG-producing Pseudomonads within three different grassland soil sites and show that DAPG producers can constitute part of the *Pseudomonas* population in the range of 0.35-17% at these sites. Finally, we show that the biosensor enables detection of DAPG produced by non-*Pseudomonas* species.

Our studies show that a whole-cell biosensor for DAPG detection can facilitate isolation of bacteria that produce this important secondary metabolite and provide insight into the population dynamics of DAPG producers in natural grassland soil.

**IMPORTANCE:** The interest has grown for bacterial biocontrol agents as biosustainable alternatives to pesticides to increase crop yields. Currently, we have a broad knowledge of antimicrobial compounds, such as DAPG, produced by bacteria growing in the rhizosphere surrounding plant roots. However, compared to the rhizosphere niches, the ecological role of DAPG as well as the distribution and dynamics of DAPG-producing bacteria remains less well understood for other environments such as bulk and grassland soil. Currently, we are restricted to chemical methods with detection limits and time-consuming PCR-based and probe-hybridization approaches to detect DAPG and its respective producer. In this study, we have developed a whole-cell biosensor, which can circumvent the labor-intensive screening process, as well as increase the sensitivity at which DAPG is detected. This enables quantification of relative amounts of DAPG-producers, which in turn increases our understanding of the dynamics and ecology of these producers in natural soil environments.

## INTRODUCTION

Secondary metabolites are well-known for their potential as drugs in the medical industry. They were initially defined as dispensable, non-vital compounds to their respective producers. However, recent advances in genome mining and microbial ecology are beginning to shed light on the prevalence, role and importance of these metabolites in natural environments. In soil ecology, species of fluorescent *Pseudomonas* isolated from naturally suppressive soils have received a great deal of attention due to their production of antimicrobial secondary metabolites, such as 2,4-diacetylphloroglucinol (DAPG). Suppression of wheat take-all disease caused by *Gaeumannomyces graminis* var. *tritici* was shown to be induced by years of crop monoculture and was associated with the root colonization of DAPG-producing fluorescent *Pseudomonas* (1). Moreover, production of DAPG was shown to be involved in disease control against the causative agent of tobacco black root rot, *Thielaviopsis basicola* (2). It has also been demonstrated that DAPG has antibacterial properties against the pathogen *Erwinia carotovorum* subsp. *atroseptica* causing soft rot of potatoes (3).

The biosynthetic gene cluster related to DAPG production in *Pseudomonas* comprises eight genes, *phlACBDEFGH* (4, 5). Proteins encoded by the operon of *phlACBD* are responsible for the synthesis of DAPG (4). The type III polyketide synthase, PhlD, initially condenses three malonyl coenzyme A molecules into phloroglucinol (PG), which is further acetylated by the enzyme complex of PhlACB into monoacetylphloroglucinol (MAPG) and DAPG (6). PhlE was identified as a putative membrane transporter with similarities to a known efflux pump in *Staphylococcus aureus* (4). The protein product of *phlG* has been described as a hydrolase that catalyses the degradation of DAPG to MAPG, thus controlling the intracellular levels of DAPG (5, 7). Both *phlF* and *phlH* encode *tetR*-like repressors that inhibit transcription of *phlACBD* and *phlG*, respectively (5, 8). PhlF and PhlH bind to operator sites located in promoters upstream of the genes they regulate, thereby sterically blocking transcription. DAPG serves as the ligand for both repressors. Thus, in the presence of DAPG repression is relieved, leading to expression of *phlACBD* and *phlG*, which in turn leads to induced DAPG biosynthesis and post-translational regulation (5, 8).

The complex of species belonging to the group, *Pseudomonas fluorescens*, has been well characterized over the past three decades, due to the potential of several species to act as biocontrol agents in agriculture. A recent study surveyed the phylogenetic relationship between 166 type strains of *Pseudomonas* (among which 66 belonged to the *P. fluorescens* group) based on amino acid sequences of 100 gene orthologues, which further proposed the existence of 10 subgroups within the *P. fluorescens* clade (9). However, despite the vast diversity among *P. fluorescens*, only few species belonging to two subgroups, *P. protegens* and *P. corrugata*, are known to produce DAPG (1, 8). With the advances in genome mining and the increased availability of complete genomes, the biosynthetic gene cluster *phlACBDE* was recently identified in *Pseudomonas* species outside of the *P. fluorescens* group, as well as in two genera of β-proteobacteria (10). However, in these cases production of DAPG has not yet been demonstrated.

Identification and enumeration of DAPG-producing microorganisms has, to our knowledge, exclusively relied on DNA probe hybridization and PCR-based techniques. One of the most commonly employed techniques uses colony hybridization combined with a confirmatory PCR to verify the presence of *phlD* (11). A more recent method involves culture-independent real-time PCR to quantify populations of DAPG-producing *Pseudomonas* in the plant rhizosphere (12). A different approach is to quantify the amount of DAPG produced *in situ* by chemical analysis. Bonsall et al. demonstrated that an optimized extraction protocol enabled quantification of DAPG isolated directly from the plant rhizosphere (13). While these techniques have clear advantages, there are several drawbacks that also exist. PCR-based methods are limited by DNA binding of specific primers and measures have to be taken to address the quantity of diverse genotypes of DAPG-producers. Chemical identification, on the other hand, is restricted by detection limits, which is directly correlated to the size of the bacterial population.

In recent years, the synthetic biology toolbox has expanded rapidly and the use of genetically engineered molecular circuits to sense molecules and conditions of interest has gained increased attention. These developments have given rise to whole-cell biosensors that utilize natural regulatory systems engineered to detect metabolites and small molecules (14, 15). Whole-cell biosensors rely on molecule recognition to either activate transcription or lift repression of a reporter gene and are thus often highly sensitive with detection limits in the nano-to micromolar range (16–18). Furthermore, whole-cell biosensors are tunable by addition or alteration of genetic parts, which allows for higher sensitivity and increased specificity (15, 19). Lastly, biosensors may also be implemented as biological detectors for uncovering metabolic activities *in situ* (20, 21).

Studies of DAPG-producing *Pseudomonas* species have predominantly focused on rhizosphere niches, whereas the ecological role of DAPG as well as the distribution and dynamics of DAPG-producing bacteria is not well understood for other environments such as bulk soil and grassland. Here, we construct a whole cell biosensor as an alternative and efficient approach for detection of DAPG and directed isolation of DAPG-producing bacteria from environmental samples.

## MATERIALS AND METHODS

### Strains, media and growth conditions

Plasmid cloning and genetic circuit characterization were performed in *Escherichia coli* K12 Δ*lacIZYA* or *E. coli* CC118-*λpir*. Cells were cultured in Luria-Bertani (LB) broth (Lennox, Merck, St. Louis, MO, USA) with appropriate antibiotics. The antibiotic concentration used was 25 µg ml^−1^ for kanamycin, 10 µg ml^−1^ for chloramphenicol and 8 µg ml^−1^ for tetracycline. The engineered whole-cell biosensor was cultured by inoculating a single colony in 5 ml LB broth supplemented with kanamycin and incubating overnight at 37° C with shaking (200 rpm). For characterization of the biosensor response to DAPG-producers, control strains were routinely cultured by inoculating a single colony in 5 ml LB broth and incubating overnight at 30° C with shaking (200 rpm). Control strains include: *Pseudomonas putida* KT2440, *Pseudomonas protegens* DTU9.1 (previously isolated by our group), *Pseudomonas protegens* DTU9.1 Δ*phlACBD* (see below) and *Chromobacterium vaccinii* MWU328.

### Plasmid circuit construction

Plasmid construction and DNA manipulation was performed following standard molecular biology techniques. The strain *E. coli* K12 Δ*lacIZYA* was transformed with all plasmid constructs by chemical transformation. The plasmid pAJM847 (Accession: MH101727.1), comprising the P_lacIQ_ promoter, *phlF*, the induction operon terminator (IOT) and the P_*phlF*_ promoter, was a kind gift from Christopher Voigt (22). The genetic circuit from pAJM847 was re-organized into pSEVA225T (Accession: KC847299.1) (23) to obtain the DAPG biosensor. To this end, the fragment containing the P_lacIQ_ promoter, *phlF* and the induction operon terminator was PCR amplified with AvrII and EcoRI overhangs. Purified PCR product was digested with appropriate restriction enzymes and inserted in pSEVA225T to yield pSEVA225::P_lacIQ_*-phlF*. Subsequently, the fragment containing the P_*phlF*_ promoter was PCR amplified with EcoRI and HindIII overhangs. Purified PCR product was restriction-digested and inserted in pSEVA225::P_lacIQ_*-phlF* to yield the *lacZ* version of the DAPG biosensor (pSEVA225::DAPG_lacZ_). The *lux* operon (*luxCDABE*) was PCR amplified from pUC18-mini-TN7T-Gm-*lux* with HindIII and SpeI overhangs. Purified PCR product was restriction-digested and inserted in pSEVA225::DAPG_lacZ_ to yield the *lux* version (pSEVA226::DAPG_lux_).

### Deletion of *phlACBD* by allelic replacement

To abolish DAPG production in *P. protegens* DTU9.1, the biosynthesis genes, *phlACBD*, were deleted by allelic replacement according to Hmelo et al. (24). In short, DNA fragments directly upstream of *phlA* and directly downstream of *phlD* were PCR amplified. The fragments were joined by splicing-by-overlap extension PCR with XbaI and SacI overhangs. The purified PCR product was restriction-digested and inserted in pNJ1 (25). The resulting plasmid was mobilized into *P. protegens* DTU9.1 via triparental mating with *E. coli* HB101 harbouring the helper plasmid pRK600. Merodiploid transconjugants were initially selected on *Pseudomonas* Isolation Agar (PIA, Merck, St. Louis, MO, USA) supplemented with 50 µg ml^−1^ tetracycline. A second selection was performed on NSLB agar (10 g l^−1^ tryptone, 5 g l^−1^ yeast extract, 15 g l^−1^ Bacto agar) with 15% sucrose. Candidates for successful deletion were confirmed by PCR and verified by Sanger sequencing at Eurofins Genomics.

### Luminescence dose/response microplate assay

Overnight cultures of the whole-cell biosensor harbouring pSEVA226-DAPG_lux_ were prepared in six biological replicates as described above. 96-well black, clear bottom microplates (In Vitro, Denmark) were prepared with LB broth supplemented with kanamycin and varying concentrations of PG, MAPG or DAPG (0, 0.005, 0.01, 0.02, 0.039, 0.078, 0.156, 0.3125, 0.625, 1.25, 2.5, 5, 7.5, 10 and 15 µM). The overnight cultures were diluted to inoculate the microplates to an initial OD_600_ = 0.01. The plates were sealed with a semipermeable membrane (Breathe-Easy, Merck, St. Louis, MO, USA) and incubated in a Cytation5 microplate reader for 8 hours at 37° C with shaking (600 rpm) with continuous measurements of luminescence and absorbance at OD_600_. This data and any other microplate assay data was collected with the Gen5 2.07 software and exported to Excel 2016 and GraphPad for data analysis.

### β-galactosidase dose/response microplate assay

Overnight cultures of the whole-cell biosensor harbouring pSEVA225-DAPG_lacZ_ were prepared in triplicate as described above. Transparent 96-well microplates (TPP, Merck, St. Louis, MO, USA) were prepared with LB broth supplemented with kanamycin and varying concentrations of PG, MAPG or DAPG (0, 0.625, 1.25, 2.5, 5, 7.5, 10 and 15 µM). The overnight cultures were diluted to inoculate the microplates to an initial OD_600_ = 0.01. Cultures were grown for 3 hours at 37° C with shaking (600 rpm), followed by measuring the end-point absorbance at OD_600_. Subsequently, 20 µl from each well was transferred to new transparent 96-well microplates and mixed with 80 µl permeabilization buffer (0.1 M Na_2_HPO_4_, 0.02 M KCl, 0.002 M MgSO_4_, 0.8 mg ml^−1^ hexadecyltrimethylammonium bromide, 0.4 mg ml^−1^ sodium deoxycholate, 5.4 µl ml^−1^ β-mercaptoethanol) (26). The plates were incubated at 30° C with shaking (600 rpm) for 30 minutes to facilitate cell lysis. Next, 28 µl lysed cell culture from each well was transferred to 96-well black microplates and mixed with 172 µl substrate solution (0.06 M Na_2_HPO_4_, 0.04 M NaH_2_PO4, 1 mg ml^−1^ o-nitro-phenyl-β-D-galactopyranoside, 2.7 µl ml^−1^ β-mercaptoethanol) (26). The plates were sealed with a semipermeable membrane (Breathe-Easy, Merck, St. Louis, MO, USA) and incubated in a Cytation5 microplate reader for 16 hours at 37° C with shaking (600 rpm) with continuous measurements of absorbance at OD_420_. Data was collected and analysed, as mentioned above.

### Detecting DAPG from bacterial cultures grown on agar surfaces

Overnight cultures of the whole-cell biosensor harbouring pSEVA225-DAPG_lacZ_ and the control strains *P. putida* KT2440, *P. protegens* DTU9.1, *P. protegens* DTU9.1 Δ*phlACBD* and *C. vaccinii* MWU328 were prepared as described above. The biosensor was normalized to OD_600_ = 1.0 and spread on King’s agar B supplemented with malt extract (KBmalt) (20 g l^−^ 1 Proteose peptone No. 3, 1.5 g l^−1^ K_2_HPO_4_, 1.5 g l^−1^ MgSO_4_, 7.5 g l^−1^ malt extract, 10 ml l^1^ glycerol and 20 g l^−1^ Bacto agar) supplemented with 25 µg ml^−1^ 5-bromo-4-chloro-3-indolyl-β-D-galactopyranoside (X-gal, Thermo Fisher Scientific). Overnight cultures of the control strains were normalized to OD_600_ = 1.0 and inoculated as 20 µl spots on the agar plates containing the biosensor. Agar plates were incubated at 30° C for 24-96 hours. Plates were inspected for blue halos surrounding the bacterial spots every 24 hours.

### Detection of DAPG by LCMS

Overnight cultures of the control strains *P. putida* KT2440, *P. protegens* DTU9.1, *P. protegens* DTU9.1 Δ*phlACBD* and *C. vaccinii* MWU328 were prepared as described above. The cultures were normalized to OD_600_ = 1.0, inoculated as 20 µl spots on KBmalt and malt agar plates and incubated at 30° C for 24 hours. An agar plug (6 mm diameter) of the bacterial culture was transferred to a vial and extracted with 1 ml of isopropanol:ethyl acetate (1:3, v/v), containing 1% formic acid, under ultrasonication for 60 min. The extracts were then transferred to new vials, evaporated under N_2_, and re-dissolved in 200 µl of methanol for further sonication over 15 min. After centrifugation at 13400 rpm for 3 min, the supernatants were transferred to HPLC vials and subjected to ultrahigh-performance liquid chromatography-high resolution electrospray ionization mass spectrometry (UHPLC-HRESIMS) analysis. UHPLC-HRESIMS was performed on an Agilent Infinity 1290 UHPLC system equipped with a diode array (DAD) detector. UV–visible spectra were recorded from 190 to 640 nm. Liquid chromatography of 1 µl extract was carried out using an Agilent Poroshell 120 phenyl-hexyl column (2.1 × 150 mm, 1.9 µm) at 60° C using acetonitrile and H_2_O, both containing 0.02 M formic acid, as mobile phases. Initially, a linear gradient of 10% acetonitrile/H_2_O to 100% acetonitrile over 10 minutes was employed, followed by isocratic wash of 100% acetonitrile for 2 minutes. The gradient was returned to 10% acetonitrile/H_2_O in 0.1 minute and finally isocratic condition of 10% acetonitrile/H_2_O for 1.9 minutes, all at a flow rate of 0.35 ml min^−1^. Mass-spectrometry detection was performed in positive ionization on an Agilent 6545 QTOF MS equipped with an Agilent Dual Jet Stream electrospray ion source with a drying gas temperature of 250° C, drying gas flow of 8 l min^−1^, sheath gas temperature of 300° C and sheath gas flow of 12 L min^−1^. Capillary voltage was set to 4000 V and nozzle voltage to 500 V. Mass-spec data analysis and processing were performed using Agilent MassHunter Qualitative Analysis B.07.00.

### Isolation of fluorescent *Pseudomonas* from grassland soil

Three sites of undisturbed grassland were chosen (P5, 55° 78’88” N, 12° 55’83” E; P8, 55° 79’52” N, 12° 58’06” E; P9, 55° 79’12” N, 12° 57’51” E). Soil was collected approximately 10 centimetres below the grass surface. Five grams of soil were suspended in 30 ml of sterile water and shaken vigorously for 1 min on a Vortex mixer. The samples were subsequently serially diluted and plated onto ¼ KB (7.5 g l^−1^ King’s agar B, 10 ml l^−1^ glycerol, 7.5 g l^−1^ Bacto agar) supplemented with 100 µg ml^−1^ cycloheximide, 13 µg ml^−1^ chloramphenicol and 40 µg ml^−1^ ampicillin. Agar plates were incubated at 30° C for 48 hours. Fluorescent colonies were identified under UV light and re-streaked on LB agar plates. Species identification of the soil isolates was performed by PCR, amplifying part of the *rpoD* gene with primers (PsEG30F: 5’-ATYGAAATCGCCAARCG, PsEG790R: 5’-CGGTTGATKTCCTTGA) (27). PCR products were purified, sequenced and aligned to a database of 166 known type strains of *Pseudomonas* (9).

### Biosensor-guided identification of DAPG-producers from grassland soil

Thirty fluorescent *Pseudomonas* were randomly selected from sample site P5 both in 2018 and 2019 as described above. Isolates were cultured overnight in LB broth at 30° C with shaking (200 rpm). An overnight culture of the biosensor harbouring pSEVA225-DAPG_lacZ_ was normalized to OD_600_ = 1.0 and spread on KBmalt plates supplemented with 25 µg ml^−1^ X-gal. Overnight cultures of the *Pseudomonas* isolates were inoculated on the agar plates as 20 µl spots. Plates were incubated at 30° C for 24-48 hours. Plates were inspected for blue halos surrounding the bacterial spots every 24 hours. The DAPG-producing isolates were identified by PCR-based species identification, as described above. The phylogenetic relationship between the DAPG-producing isolates was determined by analysing a phylogenetic tree with representatives of each *P. fluorescens* subgroup (9). In short, the PCR amplified part of the *rpoD* genes were Sanger sequenced and aligned using the MUSCLE algorithm, followed by construction of a bootstrap consensus tree (500 replicates) by the neighbour-joining method in MEGA X (28).

### High-throughput screening for DAPG-producers in grassland soil

In 2019, 288 fluorescent *Pseudomonas* were randomly selected from three sample sites (P5’, 55° 78’78” N, 12° 56’07” E; P8 and P9 – see coordinates above). Fluorescent colonies were streaked on LB agar OmniTray™ (Nunc™, Nalge Nunc International, Rochester, NY, USA) and incubated for 24 hours at 30° C. Isolates were cultured in transparent 96-well microplates in terrific broth (TB; 12 g l^−1^ tryptone, 24 g l^−1^ yeast extract, 0.17 M KH_2_PO_4_, 0.72 M K_2_HPO_4_ and 5 ml l^−1^ glycerol). An overnight culture of the biosensor harbouring pSEVA225-DAPG_lacZ_ was normalized to OD_600_ = 0.5 and spread on KBmalt OmniTrays supplemented with 50 µg ml^−1^ X-gal. The *Pseudomonas* isolates were inoculated on the OmniTrays with sterile replicators. The OmniTrays were incubated at 30° C for 48 hours and inspected for blue halos surrounding the *Pseudomonas* colonies. Candidate isolates exhibiting a blue halo were screened with PCR for the presence of *phlD* with primers (B2BF: 5’-ACCCACCGCAGCATCGTTTATGAGC, BPR4: 5’-CCGCCGGTATGGAAGATGAAAAAGTC) (29). Moreover, *rpoD* was amplified from candidate colonies for species identification with primers PsEG30F and PsEG790R.

## RESULTS

### Construction of whole-cell DAPG biosensors with high sensitivity and specificity

Two whole-cell biosensors were constructed to enable specific detection of DAPG and identification of DAPG-producing bacteria. Both sensors contain an identical module for DAPG sensing in combination with either the *lux* operon or the *lacZ* gene as reporters (Figure 1A). The biosensor plasmids were constructed as repressor-mediated modules in an *E. coli* K12 Δ*lacIZYA* host. In the absence of DAPG, the TetR-like repressor protein, PhlF, binds as a dimer to the *phlO* operator site (8) in the promoter upstream of the reporter gene. As bioavailable DAPG diffuses into the cytoplasm and binds to PhlF, the repression on the target P_phlF_ promoter is relieved. Two reporter modules were chosen. The *lux* operon was used as the output reporter to obtain a highly sensitive response measured in bioluminescence units. To enable agar plate screenings for investigating the distribution and dynamics of DAPG-producing bacteria in natural microbial communities, the *lacZ* gene was used as the second output reporter. To ensure stable inhibition of the P_phlF_ promoter under non-induced conditions, the *phlF* gene is constitutively expressed from the P_lacIQ_ promoter. Both genetic circuits were introduced into a pSEVA plasmid background (Figure 1B) to allow for rapid and simple cloning, as well as efficient mobilization into distinct hosts by triparental mating. To address the sensitivity and specificity of the whole-cell biosensors, microtiter bioassays were conducted. For characterization of the *lux* version (Figure 1C), the *E. coli* host with pSEVA226-DAPG_lux_ was grown in LB broth containing varying concentrations of PG, MAPG or DAPG with continuous measurements of luminescence and cell density. PG and MAPG are both precursors of DAPG and thus similar compounds, allowing determination of biosensor specificity towards DAPG. Each data point represents the average luminescence per OD_600_ after 175 minutes of growth, which corresponds to late exponential phase (Supplementary-S1). The whole-cell biosensor exhibited excellent sensitivity towards DAPG with a response to 20 nM being statistically significantly higher than the negative control without added DAPG (Student’s t-test, *p* = 0.01). The biosensor did not respond to the concentrations of PG tested, but a minor response to MAPG at >1.25 µM was observed. For characterization of the *lacZ* version (Figure 1D), the *E. coli* host with pSEVA225::DAPG_*lacZ*_ was grown in LB broth containing varying concentrations of either PG, MAPG or DAPG to allow for enzymatic expression. Subsequently, the biosensor response was determined in a β-galactosidase microtiter assay. Enzymatic activity of transcribed *lacZ* was estimated by continuously measuring the increase in o-nitrophenol concentration over time (Supplementary-S2). The output is displayed in Miller Units (MU = (5000 · OD_420_/min) / OD_600_). Exposing the biosensor to 0.625 µM of DAPG yielded a statistically significantly higher response compared to the negative control without added DAPG (Student’s t-test, *p* = 0.0032).

**Figure 1.**
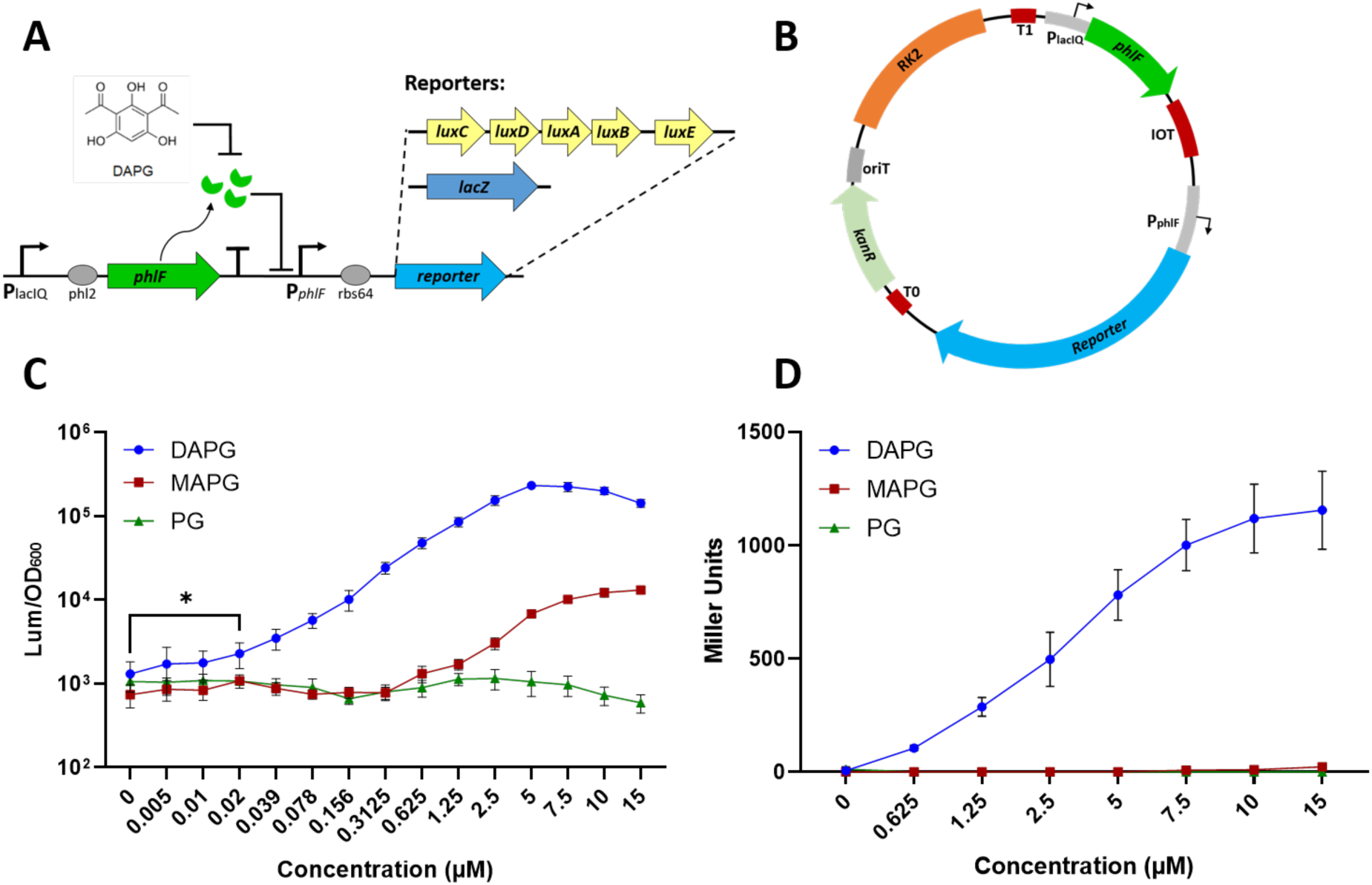
Design of highly specific and sensitive whole-cell biosensors for DAPG detection. **A)** Schematic illustration of the genetic circuit representing the DAPG biosensors with varying reporter modules. **B)** Map of the biosensor present on a pSEVA plasmid background with an RK2 replicon (orange), origin of transfer (dark grey) and a kanamycin resistance gene (light green). Terminators (T0, T1 and IOT – dark red) **C)** The response of the whole-cell biosensor harbouring pSEVA226-DAPG_lux_ to DAPG and similar molecules (PG and MAPG) measured in luminescence per OD_600_. **D)** The response of the whole-cell biosensor harbouring pSEVA225-DAPG_lacZ_ to DAPG and similar molecules (PG and MAPG) measured in Miller Units.

### Detection of DAPG production from bacterial colonies during growth on agar surfaces

After addressing the sensitivity and specificity of the biosensor, we proceeded to utilize it in the identification of DAPG-producing bacteria. We investigated the response of the whole-cell biosensor when co-inoculated with known DAPG-producing bacteria commonly found in soil (Figure 2). The biosensor harbouring pSEVA225::DAPG_*lacZ*_ was grown as a lawn on KBmalt agar (30) supplemented with X-gal. Inoculation of DAPG-producing cultures of *P. protegens* CHA0 and *P. protegens* DTU9.1 on these agar plates resulted in induction of the biosensor and production of clear blue halos surrounding the colonies after 24 hours (Figure 2A). We also constructed a Δ*phlACBD* mutant strain of *P. protegens* DTU9.1 by allelic replacement, in which the DAPG biosynthesis genes were deleted (Materials and Methods). As expected, the mutant strain did not elicit a response from the biosensor (Figure 2A) and did not produce DAPG detectable by LCMS analysis (Figure 2B). We note that DAPG production in *Pseudomonas* species has been shown to be high in growth conditions containing maltose, such as the conditions used here (8, 31). In concordance with these findings, we did not observe clear blue halos when LB agar was used (data not shown). We also tested *P. putida* KT2440, which does not contain the DAPG biosynthetic gene cluster. Similar to the *P. protegens* DTU9.1 Δ*phlACBD* mutant, *P. putida* did not elicit a response in the biosensor after 24 hours of growth. However, after prolonged incubation (>72 hours) of *P. putida* KT2440, a slight blue colouring was detected surrounding the bacterial colony (Supplementary-S3).

**Figure 2.**
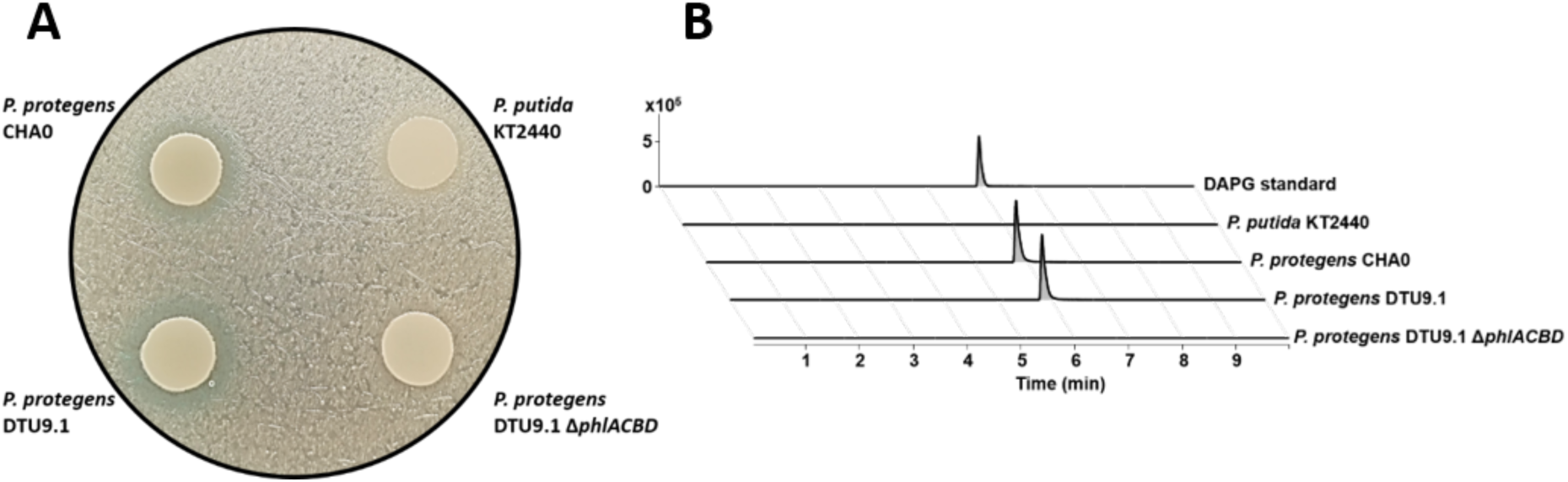
Detection of DAPG produced by bacterial colonies grown on agar surfaces. **A)** Four fluorescent *Pseudomonas* were grown on a lawn of the biosensor with pSEVA225::DAPG_*lacZ*_ on KBmalt media supplemented with X-gal. Wildtype *P. protegens* CHA0 and DTU9.1 known to produce DAPG elicited a biosensor response after 24 hours, whereas the negative controls *P. putida* KT2440 and a Δ*phlACBD* mutant of *P. protegens* DTU9.1 did not. **B)** Extracted ion chromatograms (EIC) for DAPG (*m/z* 211.0601 ± 5 ppm) of the four *Pseudomonas* extracts confirm the production of DAPG after 24 hours by *P. protegens* CHA0 and DTU9.1.

Finally, we also explored if our setup can be used to identify DAPG production in species other than *Pseudomonas*. Recently, the genome of *C. vaccinii* MWU328 was shown to contain genes with high similarity to the essential genes required for DAPG biosynthesis (*phlACBDE*) (10). We found that *C. vaccinii* MWU328 produced molecules that induced a response in the biosensor, resulting in a blue halo around the colony. A small amount of DAPG was subsequently confirmed by LCMS (Supplementary-S4).

### Biosensor-guided identification of DAPG-producing *Pseudomonas*

Next, we used our biosensor to guide the identification of DAPG-producing Pseudomonads from environmental samples. To this end, we collected soil samples from the same grassland soil site (labelled “P5”) in both August 2018 and August 2019 and randomly isolated 30 fluorescent Pseudomonas strains at both time points. This site is located in Dyrehaven, which is a Danish natural reserve, thus representing a relatively unaffected, natural soil niche. Using the approach described above, all 60 isolates were screened on KBmalt agar plates supplemented with X-gal and a lawn of the biosensor harbouring pSEVA225::DAPG_*lacZ*_. A blue halo indicative of DAPG production was observed for one isolate from 2018 (isolate P5.21) and two isolates from 2019 (isolates P5.52 and P5.53) (Figure 3A). Subsequent LCMS analysis confirmed DAPG production in all three isolates (Supplementary-S5). For taxonomic identification of the 60 isolates, part of the housekeeping gene, *rpoD*, was sequenced for each isolate. The *rpoD* sequences were aligned to a database of 165 *Pseudomonas* type strains (9). Species identification of each isolate was determined based on the highest match to the type strains using nucleotide BLAST on NCBI. The diversity of cultivable fluorescent *Pseudomonas* remained similar, as species of *P. jessenii, P. koreensis* and *P. corrugata* subgroups (as well as species of *P. putida*) were identified in the two samplings (Figure 3B). However, isolates belonging to *P. fluorescens* subgroup were only found in 2018.

**Figure 3.**
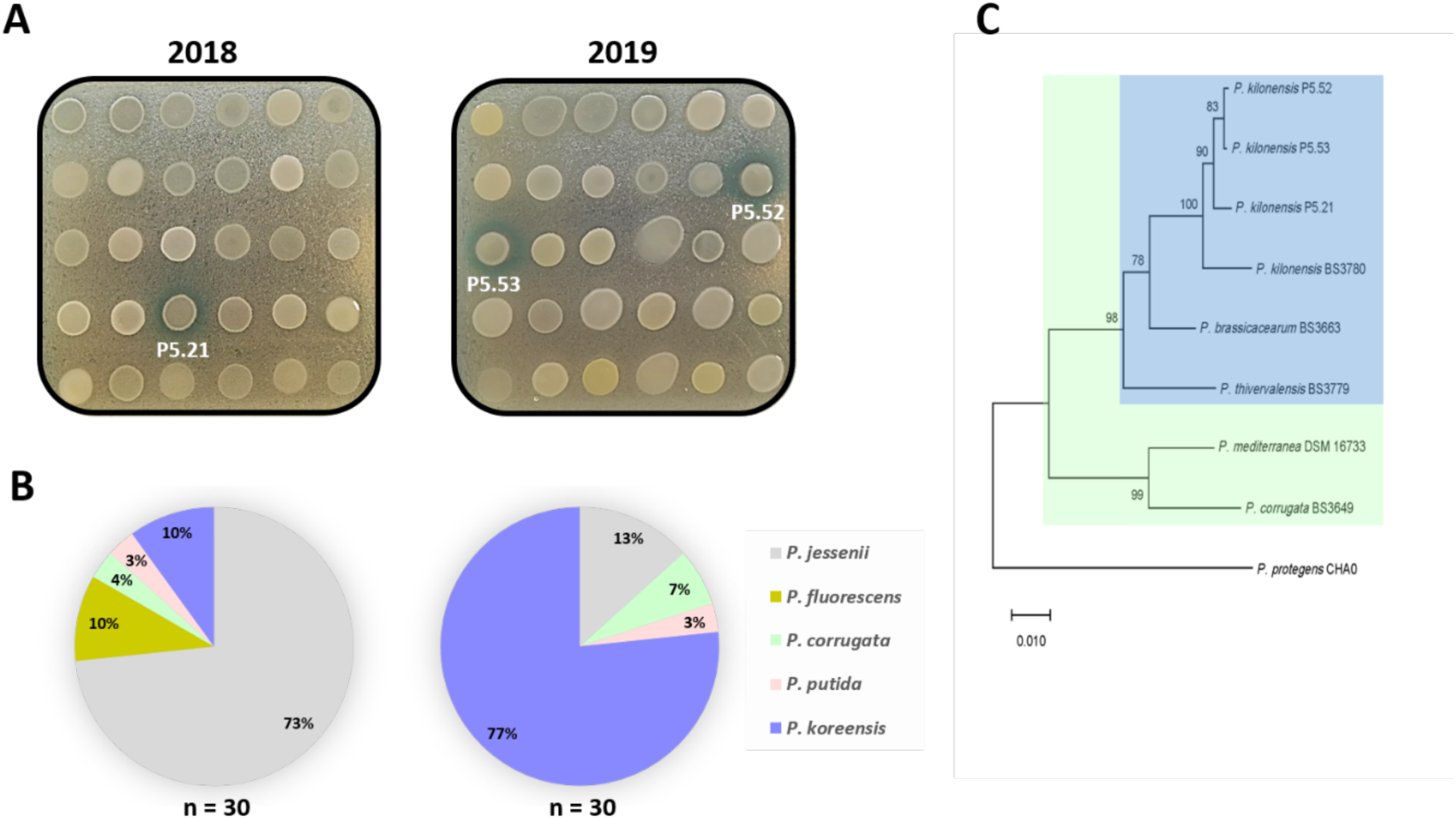
Examination of fluorescent *Pseudomonas* from grassland soil for DAPG production. **A)** 30 fluorescent *Pseudomonas* isolated from grassland soil in 2018/2019 were grown on a lawn of the biosensor with pSEVA225::DAPG_*lacZ*_ on KBmalt media supplemented with X-gal. The biosensor responded to one isolate from 2018 and two from 2019. **B)** Part of the *rpoD* gene was sequenced and aligned to a database of 165 type strains of *Pseudomonas*. The three isolates eliciting a response from the biosensor were identified as *P. kilonensis*, which is part of the *P. corrugata* subgroup (9). **C)** An *rpoD*-based Neighbor-Joining tree representing the phylogenetic relationship of the three *P. kilonensis* isolates to the *P. corrugata* subgroup. *P. protegens* CHA0 was included as outlier. A bootstrap consensus tree (500 replicates) of the *rpoD* PCR products was constructed via the Neighbor-Joining method. The bootstrap percentage values are depicted next to each branching point. The *P. corrugata* subgroup is depicted in a green box, whereas the species of this subgroup known to produce DAPG are placed in a blue box, along with the three soil isolates.

The three DAPG-producing *Pseudomonas* were identified as *P. kilonensis*. This species is part of the *P. corrugata* subgroup of fluorescent *Pseudomonas* (Figure 3C, green box). Members of this subgroup are known to harbour the biosynthetic gene cluster required for DAPG production (10). To determine the phylogenetic relationship of the three *P. kilonensis* isolates compared to the *P. corrugata* subgroup, we constructed an *rpoD*-based bootstrap consensus tree (500 replicates) with the Neighbor-Joining method (Figure 3C). As expected, the three *P. kilonensis* isolates cluster together with the members of the *P. corrugata* subgroup known to produce DAPG (marked with a blue box). Taken together, these results show that genetically highly related DAPG-producing *Pseudomonas* can be isolated from the same grassland soil site over a 12 month period.

### Measuring the frequency of DAPG-producing Pseudomonads in grassland soils

To further explore the populations and frequencies of DAPG-producing Pseudomonads in grassland soils, we sampled soil from three additional grassland sites (an area close to P5 labelled P5’, P8 and P9) (Materials and methods). In total, we isolated 288 *Pseudomonas* strains as libraries in 96-well microplates from each of the three sites (n = 864). The three libraries were then screened for potential DAPG-producers by replica-plating them onto a KBmalt agar supplemented with X-gal and a lawn of the whole-cell biosensor harbouring pSEVA225::DAPG_*lacZ*_. Simultaneously, the libraries were screened on separate plates for production of natural β-galactosidases, and isolates displaying a response were discarded from further analyses. Note that in this experimental setup (using replica-plating), the development of blue halos around DAPG-producing colonies took longer time than when larger aliquots of cultures were spotted on the agar plates. After 48 hours of incubation the biosensor elicited a response to six isolates from P5’, five isolates from P8 and 49 isolates from P9 (Table 1). Subsequently, we analysed the colonies displaying a blue halo for the presence of *phlD*, as well as their taxonomy by sequencing part of *rpoD* and aligning it to the database of *Pseudomonas* type strains (9). For P5’ and P8, one isolate from each site was confirmed to encode the polyketide synthase responsible for DAPG biosynthesis (Table 1) and both isolates were identified as *P. protegens* (Figure 4). For P9, we randomly chose 24 of the 49 isolates with a surrounding blue halo and all were confirmed to have *phlD*, where 22 of those isolates belonged to *P. kilonensis*, while the remaining two were identified as *P. protegens* (Figure 4).

**TABLE 1.** Frequencies of DAPG producers in natural soil microbiomes

|  | Total CFU/g soil<sup>†</sup> | Blue halo |  | *phlD* PCR verified |  |
| --- | --- | --- | --- | --- | --- |
|  |  | Count | Percentage | Count | CFU/g soil |
| **P5'** | $1.7 \cdot 10^4$ | 6 | 2.08% | 1/6 | 60 |
| **P8** | $5.7 \cdot 10^3$ | 5 | 1.74% | 1/5 | 20 |
| **P9** | $6.9 \cdot 10^3$ | 49 | 17.01% | 24/24* | $1.2 \cdot 10^3$ |

<sup>†</sup>: Total CFU of bacteria cultivable on $\frac{1}{4}$ KB<sup>+++</sup> media, which is predominantly *Pseudomonas*.
\*: In the case of P9, more isolates displayed a blue halo surrounding their colony than verified by *phlD* PCR.

**Figure 4.**
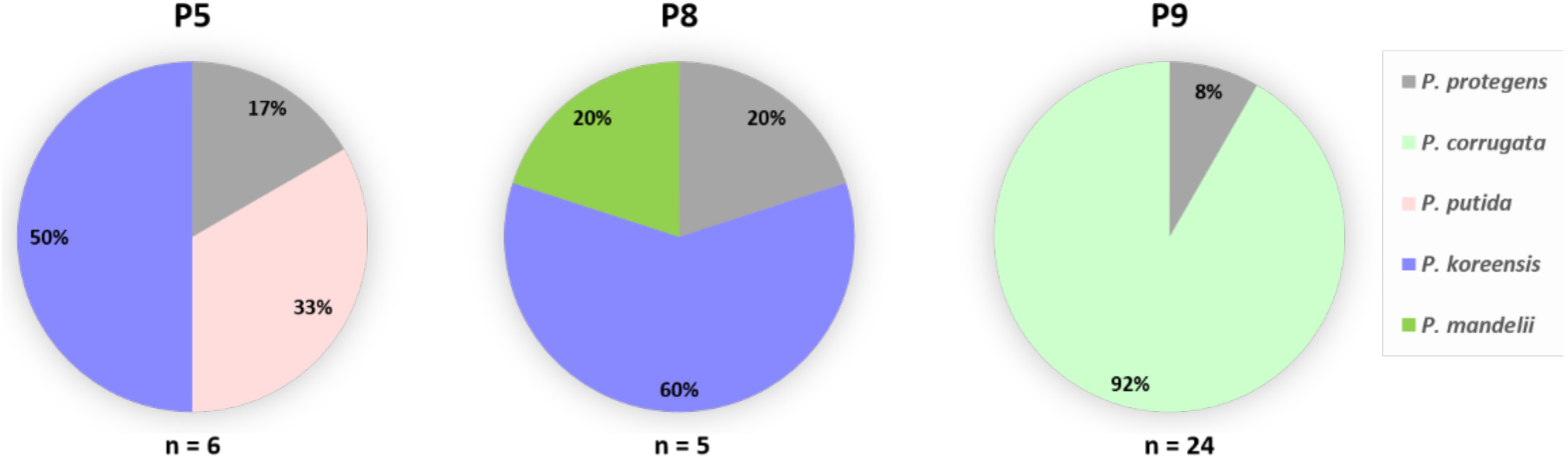
Taxonomic identification of isolates with a surrounding blue halo in the high-throughput screening assay. For each isolate, part of the *rpoD* gene was sequenced and aligned to a database of 165 type strains of *Pseudomonas*. In both P5’ and P8, one isolate was identified as *P. protegens*, whereas the remaining isolates belonged to either the *P. putida* group or *P. koreensis* and *P. mandelii* subgroups of *P. fluorescens*, according to Hesse et al. (9). In P9, the *rpoD* gene of 24/49 isolates was sequenced and the pie chart displays the distribution of the 24 isolates, with 22 identified as *P. kilonensis* and 2 as *P. protegens*.

## DISCUSSION

In this study we constructed a highly sensitive whole-cell biosensor for detection and guided isolation of DAPG-producing microorganisms. Utilization of genetic circuits to detect and report on the presence of small molecules is associated with advantages and disadvantages. Two apparent caveats associated with biosensor-guided identification are that it is only viable for cultivable organisms and it requires conditions that allow co-cultivation of the isolate of interest with the biosensor. One advantages of the biosensor is that it is not restricted to a narrow range of genotypes of DAPG-producers, which is a limiting factor of PCR-based approaches. Secondly, with a detection limit of <20 nM *in vitro* (Figure 1C) the biosensor could serve as a promising alternative to chemical identification of DAPG *in situ*, where the detection limit is in the low micromolar range (13).

We show that the biosensor is specific towards detection of DAPG. The biosensor did not respond to PG and elicited a minor response towards MAPG. This minor response was absent from its *lacZ* counterpart, which further demonstrates the sensitivity of the *lux* variant. We realize that we have only used two molecules (PG and MAPG) to represent natural DAPG analogues in our specificity assessment. It remains a possibility that other molecules can elicit a biosensor response. In a study by Yan et al. on the PhlH transcriptional regulator, it was demonstrated that multiple molecules with structural similarities to DAPG could bind to PhlH and induce a response, albeit significantly lower than the response induced by DAPG (5). Likewise, it was found that MAPG induced a minor response in the same study (5).

In order to demonstrate the biosensor response to bioavailable DAPG on agar surfaces, we inoculated DAPG producers and non-producers on top of the whole-cell biosensor on agar plates. As expected, a blue halo was observed around the DAPG producers. Additionally, a blue colouring was absent around the non-producers after 24 hours. These findings also correlated with the LCMS analysis. However, a slight blue halo was observed surrounding *P. putida* after prolonged incubation (>72 hours), suggesting that one or more compounds are being secreted by this strain during late stationary phase, which interacts with PhlF, thus relieving repression of the reporter gene.

Subsequently, the biosensor was utilized for guided identification of DAPG-producing fluorescent *Pseudomonas*. We screened 30 randomly isolated *Pseudomonas* from the P5 site in both 2018 and 2019. DAPG producers were detected in both samplings based on a clear blue halo surrounding their colonies and were identified as *P. kilonensis* species, which are known to produce DAPG (10). Part of the *rpoD* gene was sequenced for all isolates and aligned to a database of *Pseudomonas* type strains (9), which revealed a remarkably similar diversity over a 12 month period (i.e. the same *Pseudomonas* subgroups were sampled at both time points). However, despite the low sampling depth, the species abundance appears to shift from *P. jessenii* to *P. koreensis*. From an ecological point of view, it is of interest to note that the DAPG producers seem to persist over time in relatively similar quantities.

Lastly, we sought to optimize the screening assay to a high-throughput 96-well microplate format. We isolated 288 *Pseudomonas* species from each of three soil sites (P5’, P8 and P9). The sites are located in a Danish natural reserve (Dyrehaven), thus we argued that they represent pristine grassland soil niches. It is worth noting that bulk soil is an extremely harsh environment with low nutrient availability, which might explain the low amount of CFU g^−1^ of *Pseudomonas* compared to rhizosphere environments (11–13). We identified 6 and 5 isolates with blue halos around their colonies in P5’ and P8, respectively. Yet, only one isolate from each site was confirmed to encode the polyketide synthase responsible for DAPG production. Picard et al. isolated 156 *Pseudomonas* from bulk soil, but no DAPG producers were identified, although DAPG producers were isolated at a later stage from roots of maize plants grown in the same soil (32). It was speculated that DAPG-producing *Pseudomonas* are present in bulk soil in quantities <2.6 · 10^2^ CFU g^−1^, which is comparable to the findings obtained in our study of grassland soil at P5’ (6 x 10^1^ g^−1^) and P8 (2 ⨯ 10^1^ g^−1^) (Table 1). In P9, on the other hand, 17% of the isolates displayed a blue halo around their colonies and the presence of *phlD* was confirmed for 24/24 tested isolates. However, during the course of our study it became apparent that a deer-feeding site was located near the sampling site, with wheat being the main feed. This could potentially explain the high frequency of DAPG producers in P9, as DAPG-producing *Pseudomonas*, which are known to be associated with the roots of wheat (1), might have translocated into the soil surrounding the feeding site due to animal activities.

The false positive isolates from the high-throughput screening (i.e. isolates that resulted in a biosensor response without the presence of DAPG biosynthesis genes) could potentially produce compounds similar to those made by *P. putida* (as described above), which interfere with PhlF. The *phlF* gene encoded by the biosensor was cloned from *P. protegens* CHA0 (22), where it naturally functions as a transcriptional repressor of *phlACBD* (8). The false positives identified in our screen may secrete yet unknown secondary metabolites that interact with the PhlF repressor, thus inducing biosynthesis of DAPG. This finding highlights the possibility for microbe-microbe interactions *in situ* leading to induced DAPG production by adjacent non-producers. Interestingly, we found two isolates belonging to the *P. putida* group in the high-throughput screening, which may indicate that certain species of this group produce molecules that can induce expression of DAPG. Surprisingly, we also identified isolates of *P. koreensis* and *P. mandelii* that elicit a response from the biosensor, which further enhances the potential of yet unexplored microbe-microbe interactions that could be addressed in future studies.

In conclusion, this study demonstrates the use of an engineered whole-cell biosensor for guided identification of DAPG-producing microorganisms. This approach surpasses the limits of previous PCR-based and chemical identification methods, although future optimization to further increase sensitivity and reduce unexpected response to false positives might be required.

## Acknowledgements

We would like to thank Prof. Victor de Lorenzo for providing the pSEVA plasmid used as vector background for the biosensor. Secondly, we thank Prof. Christopher Voigt for providing the plasmid pAJM847. Moreover, we want to thank Pavelas Sazinas for advice on phylogenetic analysis, as well as Susanne Koefoed for technical assistance. We thank the members of the Centre for Microbial Secondary Metabolites (CeMiSt) for discussions.

## Funding

This study was funded by the Danish National Research Foundation (DNRF137) for the Centre for Microbial Secondary Metabolites.

